# Shade-induced transcription of PIF-Direct-Target Genes precedes H3K4-trimethylation chromatin modification rises

**DOI:** 10.1101/2022.01.27.478074

**Authors:** Robert H. Calderon, Jutta Dalton, Yu Zhang, Peter H. Quail

## Abstract

The phytochrome (phy)-PIF (Phytochrome Interacting Factor) sensory module perceives and transduces light signals to Direct-Target Genes (DTGs), which then drive the adaptational responses in plant growth and development, appropriate to the prevailing environment. These signals include the first exposure of etiolated seedlings to sunlight upon emergence from subterranean darkness, and the change in color of the light that is filtered through, or reflected from, neighboring vegetation (‘shade’). Previously, we identified three broad categories of rapidly signal-responsive genes: those repressed by light and conversely induced by shade; those repressed by light, but subsequently unresponsive to shade; and those responsive to shade only. Here, we investigate the potential role of epigenetic chromatin modifications in regulating these contrasting patterns of phy-PIF module-induced expression of DTGs. Using RNA-seq and ChlP-seq, time-resolved profiling of transcript and histone 3 lysine 4 trimethylation (H3K4me3) levels, respectively, we show that, whereas the initial dark-to-light transition triggers a rapid, apparently temporally-coincident decline of both parameters, the light-to-shade transition induces similarly rapid increases in transcript levels that precede increases in H3K4me3 levels. Together with other recent findings, these data raise the possibility that, rather than being causal in the shade-induced expression changes, H3K4me3 may function to buffer the rapidly fluctuating shade/light switching that is intrinsic to vegetational canopies under natural sunlight conditions.

## Introduction

All organisms must perceive, process and react to environmental cues in order to survive and pass their genetic material onto the next generation. Land plants in particular, given their sessile lifestyle, must quickly perceive these environmental signals and respond accordingly. One particularly well-studied plant signaling system is the phytochrome (phy) family of photoreceptors (phyA to phyE in Arabidopsis), a set of red (R) and far red (FR) light-absorbing chromoproteins that transduce light signals into large-scale changes in gene expression (Tepperman et al., 2001). Upon absorption of R light, the inactive form of the phy molecule (Pr) is photoconverted into the active form (Pfr) which quickly translocates from the cytoplasm to the nucleus, initiating downstream developmental programs, directed by these expression changes (Sakamoto and Nagatani, 1996).

Experimental evidence indicates that a critical link between these downstream programs and the phy molecules is a subfamily of eight bHLH transcription factors called phy-interacting factors (PIFs) (Ni et al., 1998; Huq and Quail, 2002; Monte et al., 2004; Leivar and Quail, 2011; Pham et al., 2018). The PIFs, in particular PIF1, PIF3, PIF4 and PIF5 (called the PIF quartet), form a set of partially functionally redundant proteins that bind to a consensus sequence in the upstream region of target genes, regulating their transcriptional output (Leivar et al., 2009). The PIF quartet has been shown to physically interact specifically with the Pfr form of phytochrome B (phyB), which subsequently induces phosphorylation, ubiquitination and degradation of the transcription factor (Ni et al., 2013; 2014; 2017), thereby triggering global changes in target gene expression (Leivar et al., 2009; Leivar and Quail, 2011; Pham et al., 2018). In addition to the PIF quartet, PIF6 and PIF7 have also been shown to function in phyB signaling, with PIF7 in particular serving as a key regulator of auxin biosynthesis during the shade-avoidance response (Khanna et al., 2004; Leivar et al., 2008; Li et al., 2012). The integration of several genome-wide analyses of PIF-binding and PIF-mediated transcriptional regulation (Leivar et al., 2009; Hornitschek et al., 2012; Leivar et al., 2012; Oh et al., 2012) has led to the discovery of over 300 Direct Target Genes (DTGs) that are directly, transcriptionally regulated by PIFs (Zhang et al., 2013; Pfeiffer et al., 2014).

The relative abundance of the Pfr and Pr forms of the phyB molecule, and by extension the accumulation and activity of the PIFs, is determined by the ratio of red to far-red light in the immediate environment. The active Pfr form is favored under white-light illumination where the R/FR ratio is high, whereas the inactive Pr form is favored in the dark and in conditions where the R/FR ratio is low, such as under vegetative shading (Quail et al., 1995). As a consequence of the photoreversible nature of the phyB molecule, PIF accumulation and activity is high in darkness and in the shade. The transcriptional responses of many PIF DTGs, however, do not exhibit a photoreversible pattern (Leivar et al., 2012).

In a previous study, we were able to categorize the transcriptional responses of PIF DTGs in tothree distinct patterns: those that respond during the transition from the etiolated dark-grown state to R, those that respond during the transition from white light into simulated shade or those that respond during both transitions (Leivar et al., 2012). The differential responsiveness of these three broad sets of PIF DTGs, indicates that PIF abundance is not the sole determinant of PIF DTG expression. Core components of the plant circadian oscillator have been implicated in modulating some of these changes in gene expression (Martín et al., 2018; Zhang et al., 2020). Most recently, changes in the chromatin environment have been shown to be directly involved in triggering shade-induced transcription (Willige et al., 2021).

One form of chromatin remodeling that can modulate the transcriptional output of light-regulated genes involves the enzymatic modification of histones (Fisher and Franklin, 2011; Perrella and Kaiserli, 2016; Bourbousse et al., 2019; Martínez-García and Moreno-Romero, 2020). Methylation, acetylation and/or ubiquitination of histones have all been shown to regulate transcription of light-regulated genes (Charron et al., 2009; Bourbousse et al., 2012; Liu et al., 2013). Unique histone modification patterns at the promoters of individual PIF DTGs have the potential to underly the differential responsiveness of PIF DTGs under different environmental conditions. The accumulation of one particular mark, histone 3 lysine 4 trimethylation (H3K4me3), at the transcriptional start site (TSS) of genes has long been known to strongly correlate with transcriptional activity of those genes (Bernstein et al., 2002), but the biological function of this mark remains relatively less-well defined (Fiorucci et al., 2019). Proposed roles include facilitating transcriptional elongation (Ding et al., 2012) or serving as “transcriptional memory” (Liu et al., 2014)

Here, we have refined the list of PIF DTGs by integrating previously published ChIP binding and RNA-seq data for the PIF quartet, with newly obtained RNA-seq data from both wild-type and a mutant lacking six of the PIFs (PIF1, 3, 4, 5, 6 and 7). Using this system, we have explored the potential role of the epigenetic mark H3K4me3 in mediating the observed differential patterns of expression of PIF DTGs. Our data suggest a possible functional role for H3K4me3 in stabilizing the expression levels of DTGs in established green plants, against the rapidly switching light/shade transitions that occur naturally in leaf canopies.

## Results

### Characterization of *pifqpif6pif7* sextuple mutant

The *pif1pif3pif4pif5* quadruple mutant (hereafter *pifq*) displays a constitutively photomorphogenic phenotype when grown in darkness, indicating that these four PIFs are necessary and sufficient to control de-etiolation in response to R (Leivar et al., 2008; Leivar et al., 2009). The *pifq* mutant does not, however, exhibit a complete lack of responsiveness to simulated shade (**Figure 1**), supporting the hypothesis that additional factors are required for the complete shade avoidance response (Leivar et al., 2012). PIF7 has been implicated in playing a major role in regulating this process (Li et al., 2012; de Wit et al., 2015; Mizuno et al., 2015) with the quintuple *pifqpif7* mutant reported to show no statistically-significant shade avoidance response (Zhang et al., 2020).

**Figure 1.**
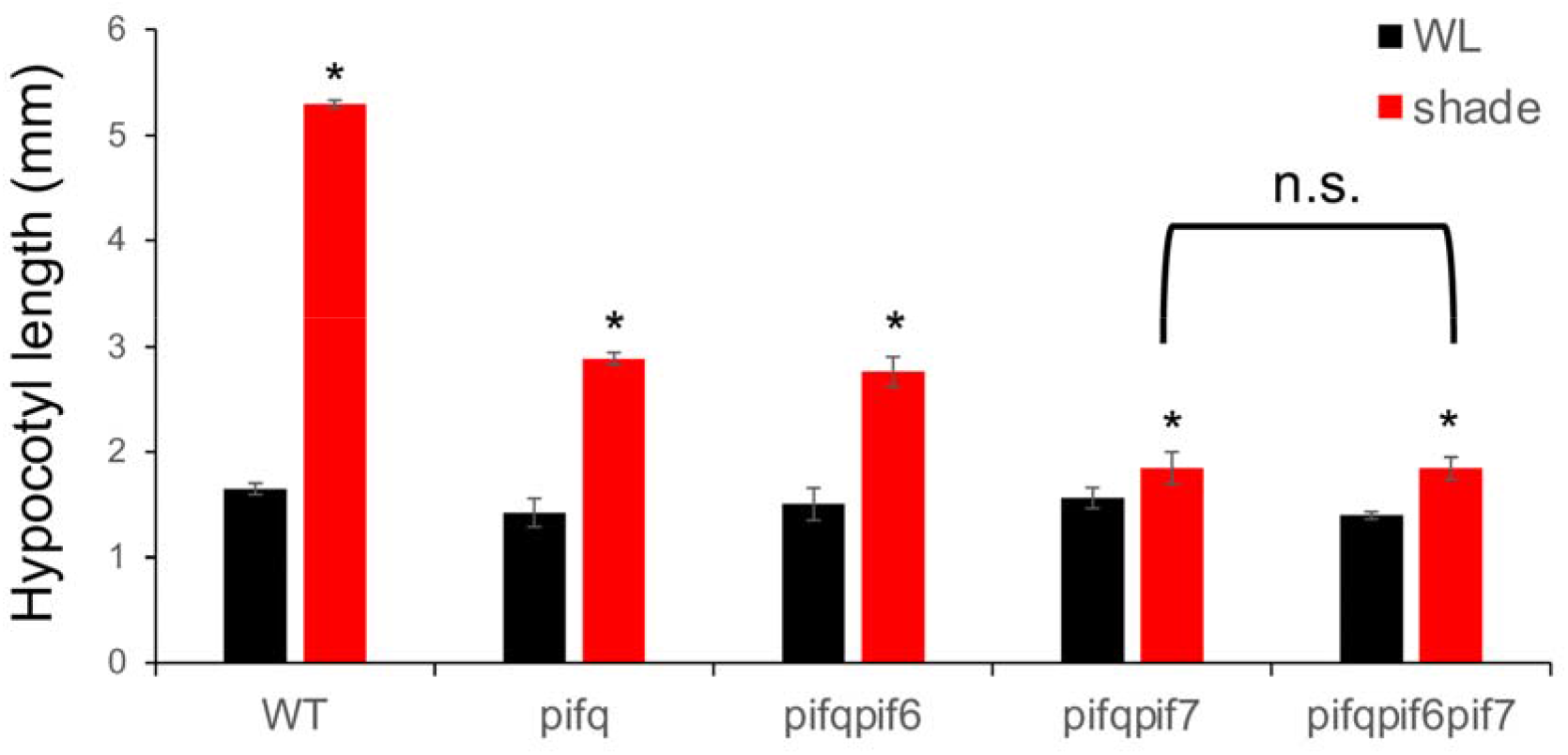
Phenotypic analysis of higher-order *pif* mutants in response to simulated shade. Hypocotyl lengths of wild-type (WT), *pifq*, *pifqpif6*, *pifqpif7* and *pifqpif6pif7* mutants grown for 6 days in white light (WL) or 2 days in WL followed by 4 days in simulated shade (shade). Data represent the mean and SE from 3 biological replicates of 30 seedlings per genotype. Asterisks indicate that the hypocotyl lengths of shade-treated seedlings are statistically significantly different from the corresponding WLc controls by Student’s t test (P < 0.05). n.s. indicates “not significantly different” (P > 0.99).

However, when we measured the shade avoidance response in the *pifqpif7* mutant under slightly different conditions to Zhang et al. (Zhang et al., 2020), we were still able to detect a small, yet statistically-significant (p < 0.05) residual shade avoidance response (**Figure 1**). A possible reason for this small difference is that the results presented here were obtained on 2-day-old seedlings exposed to simulated shade, whereas our previous experiments were performed on 3-day-old seedlings exposed to simulated shade. Alternatively, this minor residual shade-avoidance response observed under our conditions could be due to the presence of yet other members of the PIF-subfamily, such as PIF8 or PIL1 (PIF2) (Leivar and Quail, 2011; Pham et al., 2018), or to other light-responsive transcription factors. Nevertheless, we then tested whether PIF6 might be responsible for this residual response by generating a sextuple *pifqpif6pif7* (*pifS*) mutant and measuring its hypocotyl length in response to simulated shade. This sextuple mutant displayed significantly shorter hypocotyls than the wild-type in response to shade, but no significant decrease relative to the *pifqpif7* quintuple mutant (**Figure 1**). These results suggest that PIF6 plays no significant role in mediating the shade-avoidance response, consistent with its proposed role in seed dormancy and development (Penfield et al., 2010).

### Generation of a high-confidence list of PIF DTGs and subcategorization into E, ES and S classes

Many PIF direct target genes (DTGs) have been previously observed to be upregulated in the presence of the PIFs while others are downregulated. For the purposes of this study, we focused only on PIF-induced genes (*i.e*. those genes which appear to require the PIFs for high levels of transcription) because PIFs have been shown to have intrinsic activating activity (Huq et al., 2004; Al-Sady et al., 2008; de Lucas et al., 2008; Dalton et al., 2016).

In brief, we first integrated the data from a previously published RNA-seq experiment on dark-grown seedlings exposed to 1h of R light (Pfeiffer et al, 2014) with a new RNA-seq time-course experiment of white light (WL)-grown seedlings exposed to 3h of simulated shade (shade-light). We then combined previously published RNA-seq data from the *pifq* mutant grown in darkness (Pfeiffer et al., 2014), with new RNA-seq data, that were obtained using the *pifqpif6pif7* mutant (*pifS*) grown in WL and exposed to 3h shade-light. Lastly, we used previously published data to identify those genes whose promoters were found to be bound by PIF1, PIF3, PIF4, PIF5 and/or PIF7 (no genome-wide binding data are available for PIF6) (Hornitschek et al., 2012; Oh et al., 2012; Zhang et al., 2013; Pfeiffer et al., 2014; Chung et al., 2020). By selecting only the genes that met all three of our criteria (light-responsiveness, PIF-dependence and PIF-binding), we obtained 169 candidate PIF-induced, red-light repressed and/or shade-light-induced DTGs **(Table 1**).

**Table 1.**
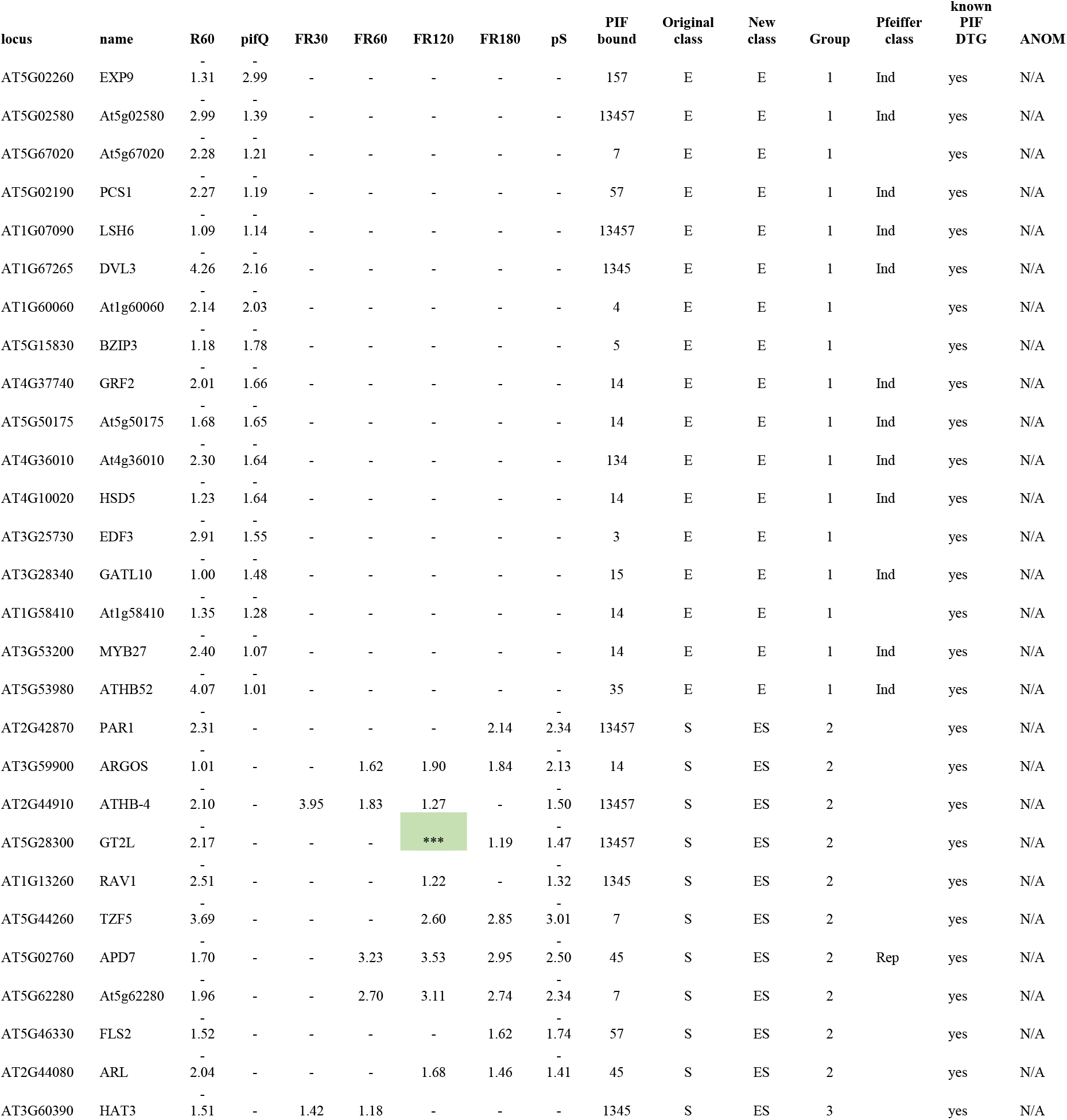
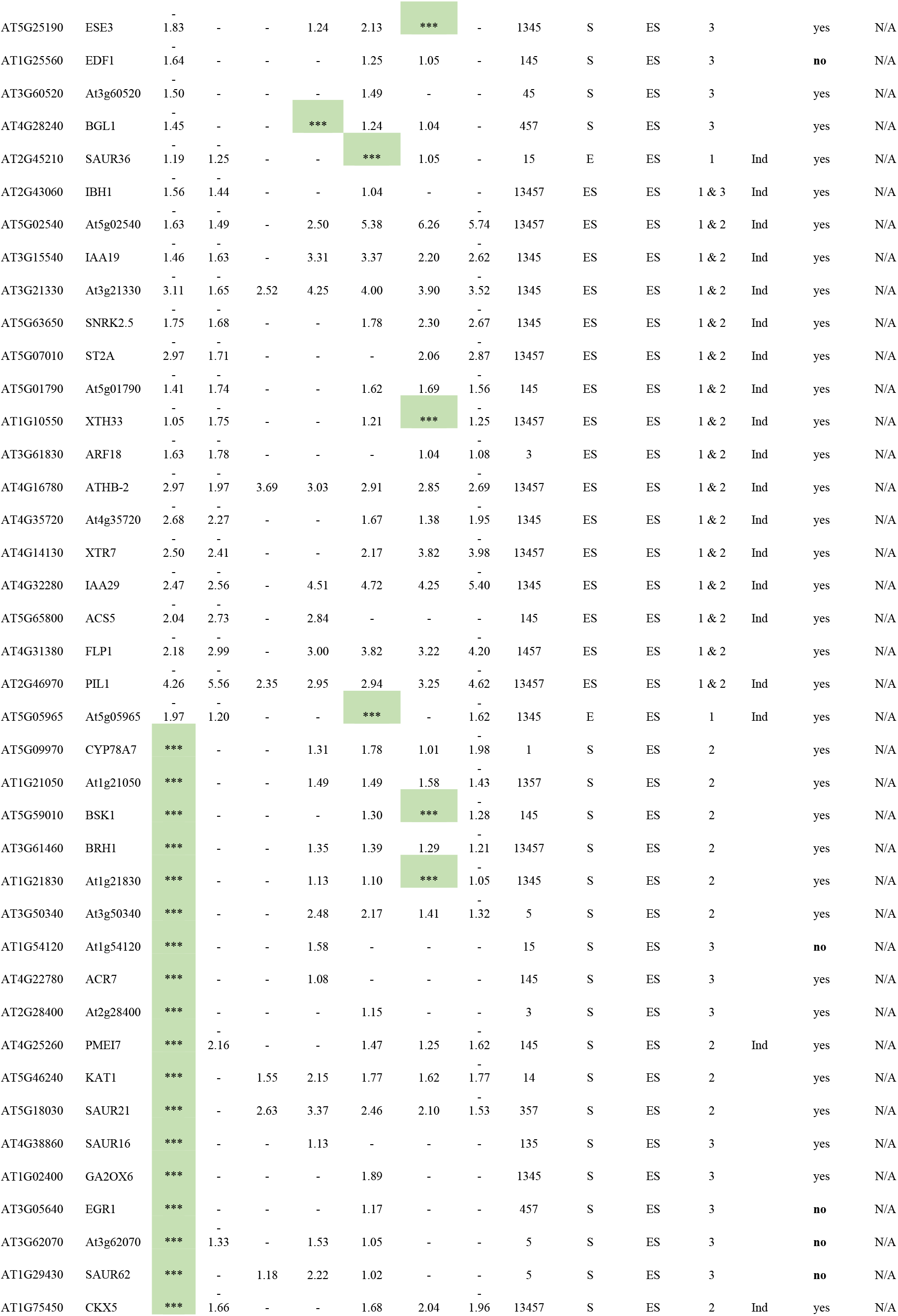
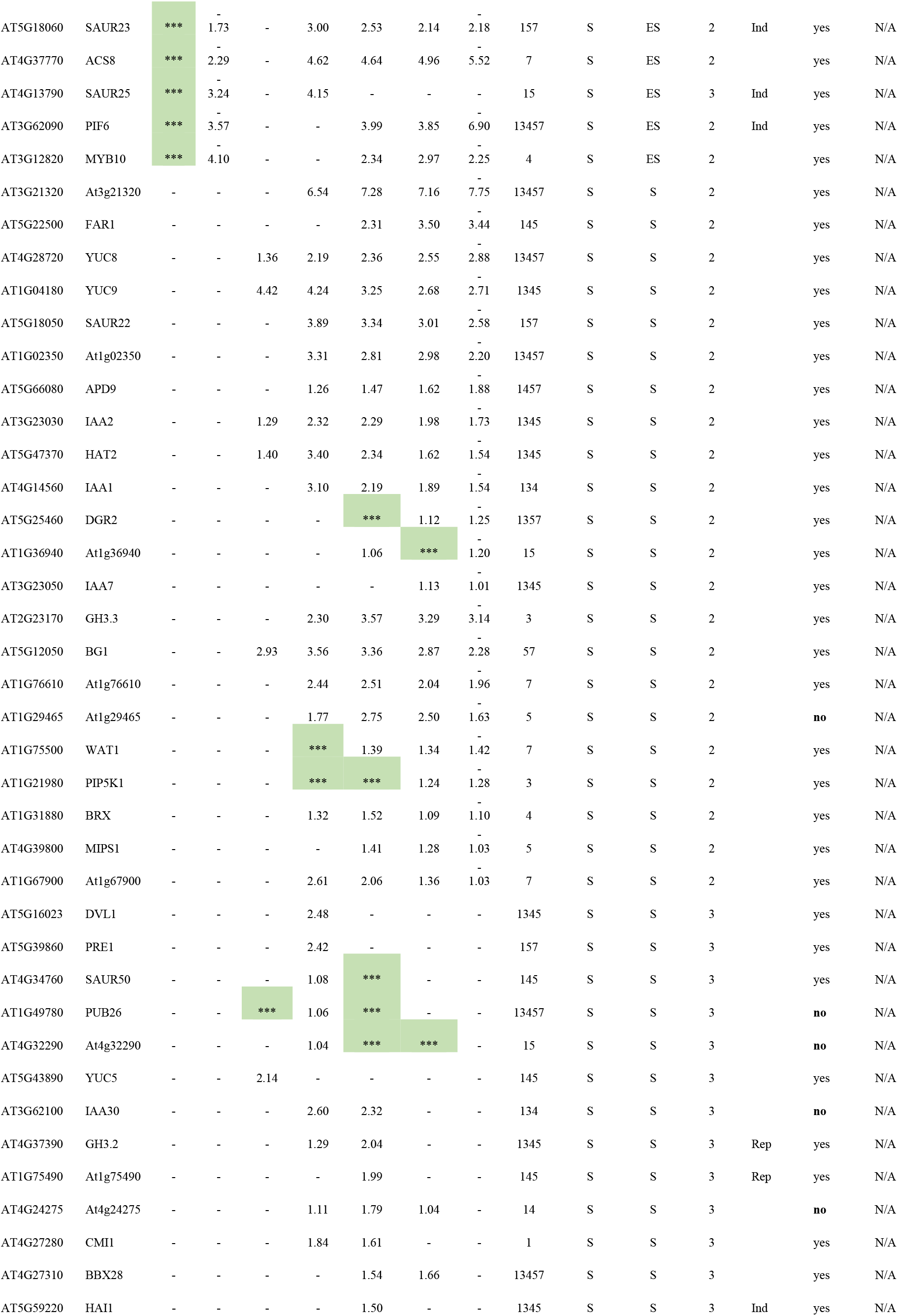
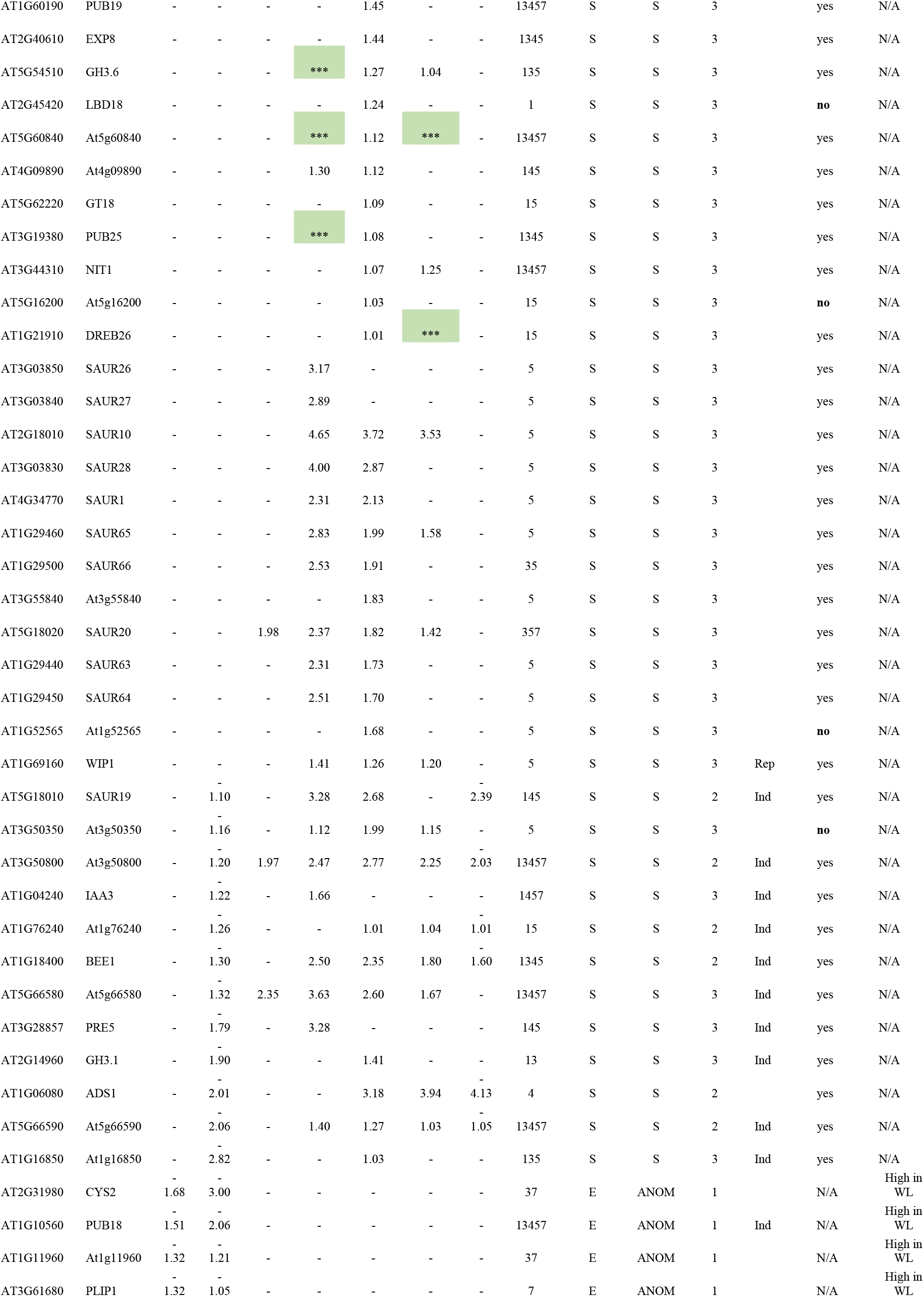
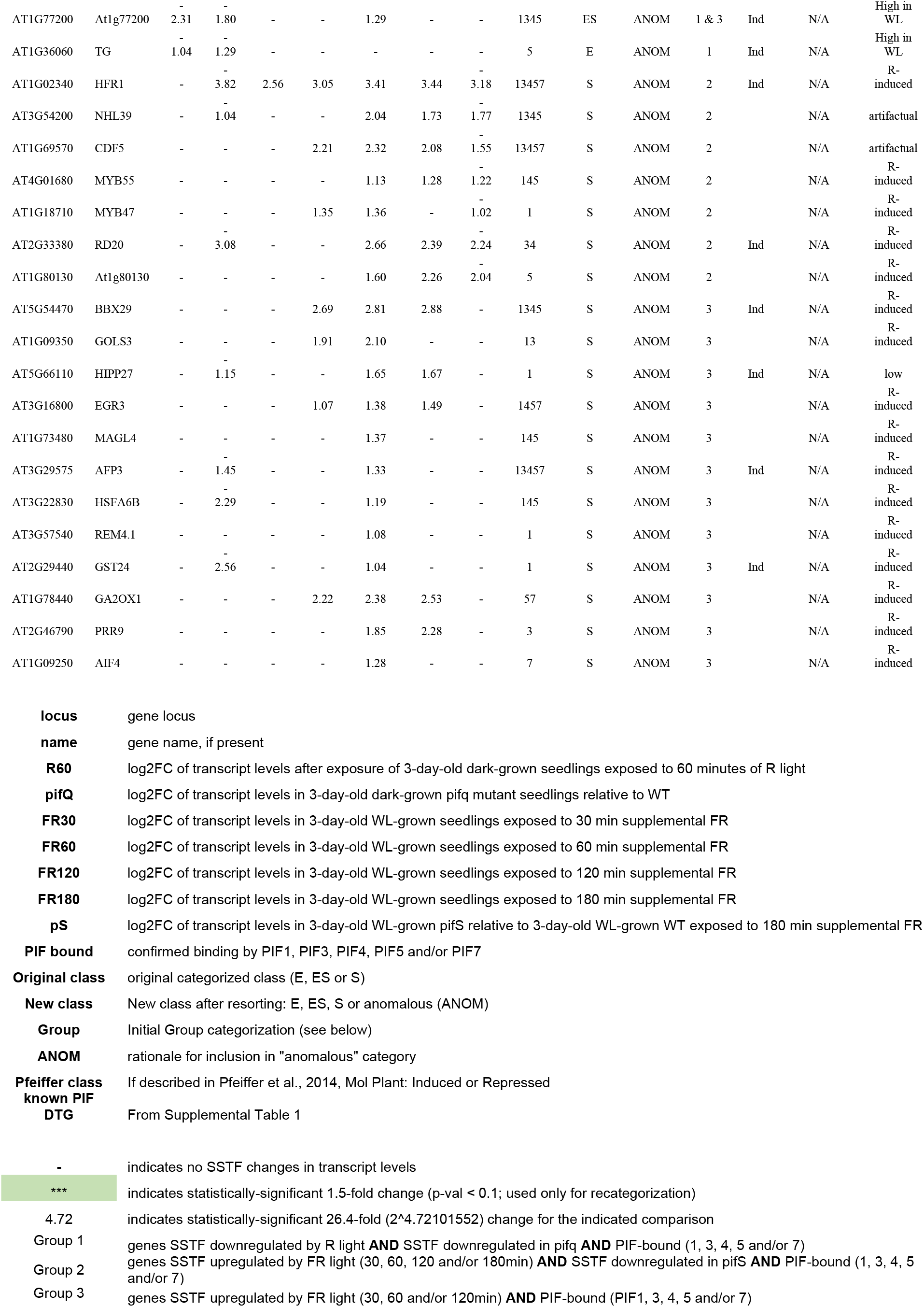
List of all candidate PIF DTGs and categorization into etiolation (E), shade (S) or etiolation and shade (ES) responsive genes.

As described in Leivar *et al*. (Leivar et al., 2012), PIF DTGs may be broadly classified into one of three classes: re-labeled here as E, ES and S (E for **E**tiolation-induced only; ES for both **E**tiolation- and **S**hade-induced; and S for **S**hade-induced only) (**Figure 2)**. We therefore subdivided our combined 169 shade-light-induced and red-repressed PIF DTGs into these classes based on their patterns of expression during the D to R, and WL to shade-light transitions. Using these criteria, our initial list of 169 genes was found to contain 24 E genes, 17 ES genes and 128 S genes (**Table 1**). Upon further analysis, we removed 25 genes that exhibited various anomalous expression profiles and resorted the remaining 144 genes using relaxed cutoff criteria. This resulted in a redistribution between the classes so that the final numbers of genes in each class were: 17 E genes, 56 ES genes and 71 S genes **(Table 1**).

**Figure 2.**
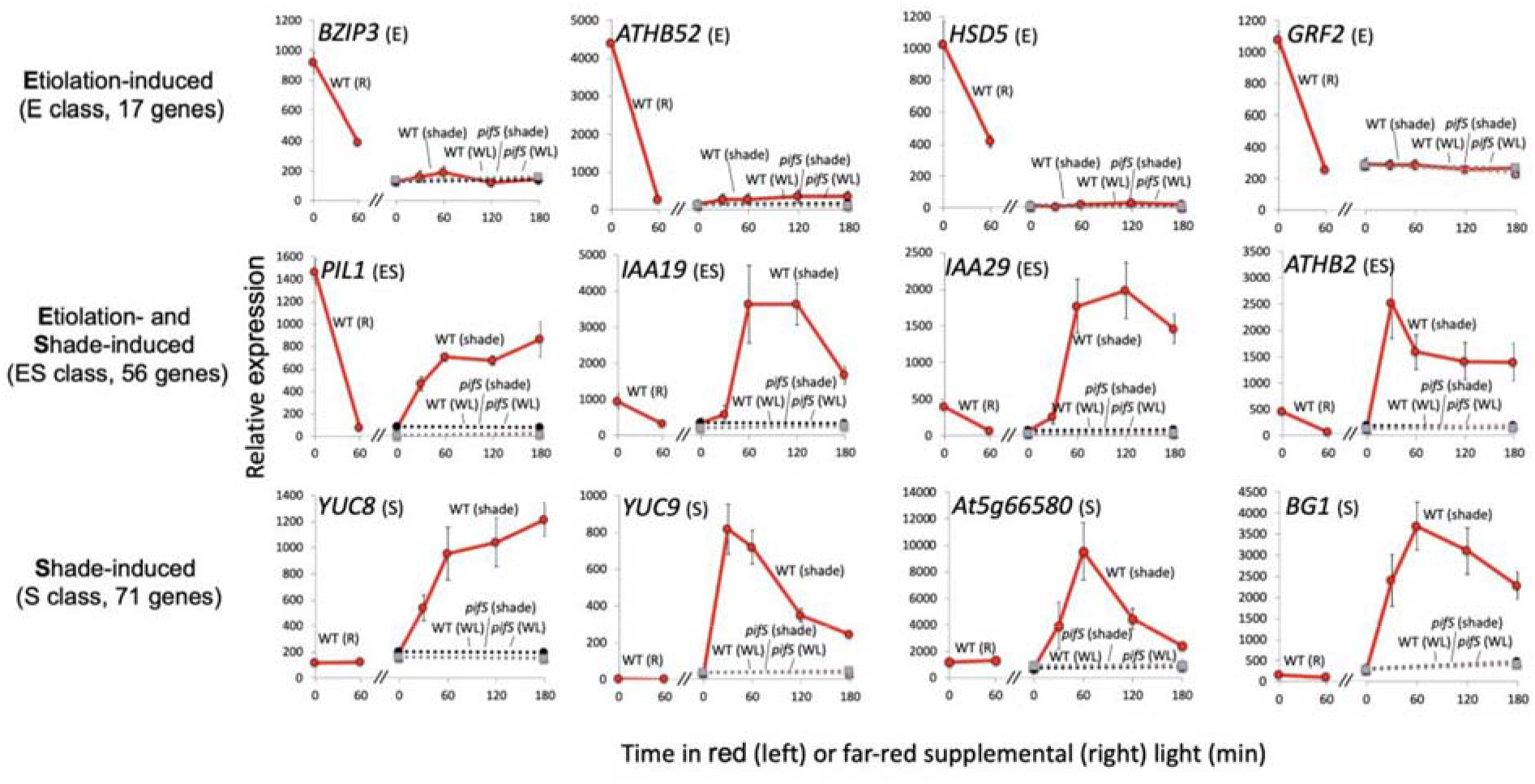
PIF-activated direct target genes (DTGs) can be subdivided into three categories based on their responses to red light and simulated shade. Examples of transcript time course profiles for Etiolation-induced only (E) genes (BZIP3, ATHB2, HSD5 and GRF2), Etiolation and Shade-induced (ES) genes (PIL1, IAA19, IAA29 and ATHB2) and Shade-induced only (S) genes (YUC8, YUC9, At5g02865 and BG1) class genes. Left subpanel shows the effect of 60 min red light on the transcript levels in 3-day-old dark-grown seedlings (wild-type, solid red line). Right subpanel shows the effect of 30, 60, 120 and 180 min of FR-enriched WL or continuous WL on transcript levels in 3-day-old WL-grown seedlings (wild-type, FR: solid red line; *pifS*, FR: dotted black line; wild-type, WL: dotted red line; *pifS*, WL: dotted gray line). Error bars indicate SE.

### Examination of potential epigenetic regulation of DTGs

We next tested our hypothesis that the variation in transcriptional responses of the PIF-activated DTGs to darkness and shade might be due to differences in histone tail modifications. One histone mark, H3K27me3, has already been linked to light-mediated transcriptional repression (Charron et al., 2009). Because we were focused on loci at which PIFs act as transcriptional activators, we sought to examine the levels of a histone mark associated with active transcription. One such mark, H3K4me3 is both correlated with actively transcribed genes (Bernstein et al., 2002) and inversely correlated with H3K27me3 levels (Zhang et al., 2009). We therefore chose to assay H3K4me3 levels at the transcriptional start sites (TSS) of E, ES and S genes by ChIP-seq. We measured H3K4me3 levels in dark-grown seedlings and in WL-grown seedlings after exposure to 0, 30, 60, 120 and 180 min of simulated shade, and after 180 min of further retention in WL. We also measured H3K4me3 levels in WL-grown *pifS* seedlings after 0 and 180 min of simulated shade and after 180 min of continued WL.

As expected, H3K4me3 levels for E class genes were higher in D than in WL and simulated shade (**Figure 3**). On average, H3K4me3 levels for ES and S class genes increase over the course of the shade treatment and this increase is attenuated in the *pifS* mutant **(Figure 4**). In both classes, however, the increase only occurs after 60 minutes of FR, while an increase in transcript level abundance is already visible after 30 minutes of FR. Both classes also exhibit a transient reduction in H3K4me3 levels after 30 minutes of FR. Collectively, these data indicate that the shade signal induces a transcriptional response prior to the induction of increased H3K4 trimethylation in these DTGs.

**Figure 3.**
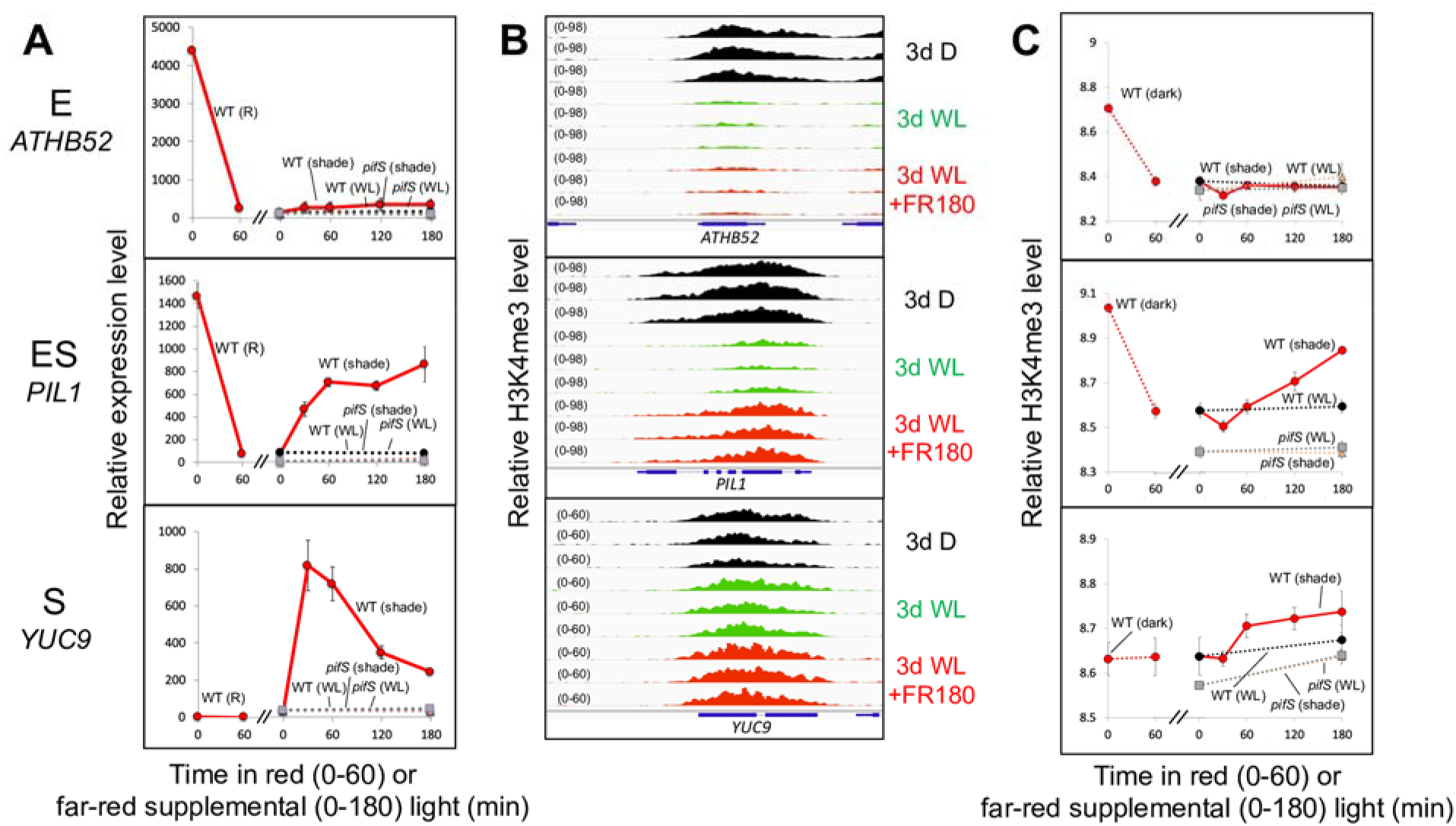
Transcript levels are broadly correlated with H3K4me3 levels for PIF DTGs belonging to Etiolation-induced only (E), Etiolation and Shade-induced (ES) and Shade-induced only (S) classes. **A)** Average relative transcript levels as measured by RNA-seq of ATHB52 (E class, top), PIL1 (ES class, middle) and YUC9 (S class, bottom). Left subpanel shows the effect of 60 min red light on the transcript levels in 3-day-old dark-grown seedlings (wild-type, solid red line). Right subpanel shows the effect of 30, 60, 120 and 180 min of FR-enriched WL or continuous WL on transcript levels in 3-day-old WL-grown seedlings (wild-type, FR: solid red line; *pifS*, FR: dotted black line; wild-type, WL: dotted red line; *pifS*, WL: dotted gray line). Error bars indicate SE.**B)** H3K4me3 enrichment as measured by ChIP-seq of ATHB52 (top), PIL1 (middle) and YUC9 (bottom) in 3-day-old dark-grown seedlings (3d D, black), 3-day-old WL-grown seedlings (3d WL, green) and 3-day-old WL-grown seedlings after 180 min of FR-enriched WL (3d WL +FR180, red). Data from each of three biological replicates are shown. **C)** Average relative H3K4me3 levels of ATHB52 (top), PIL1 (middle) and YUC9 (bottom). Left subpanel shows the levels in 3-day-old dark-grown seedlings and the levels in 3-day-old WL-grown seedlings (wild-type, connected by dashed red line). Right subpanel shows the effect of 30, 60, 120 and 180 min of FR-enriched WL or continuous WL on transcript levels in 3-day-old WL-grown seedlings (wild-type, FR: solid red line; *pifS*, FR: dotted black line; wild-type, WL: dotted red line; *pifS*, WL: dotted gray line). Error bars indicate SE.

**Figure 4.**
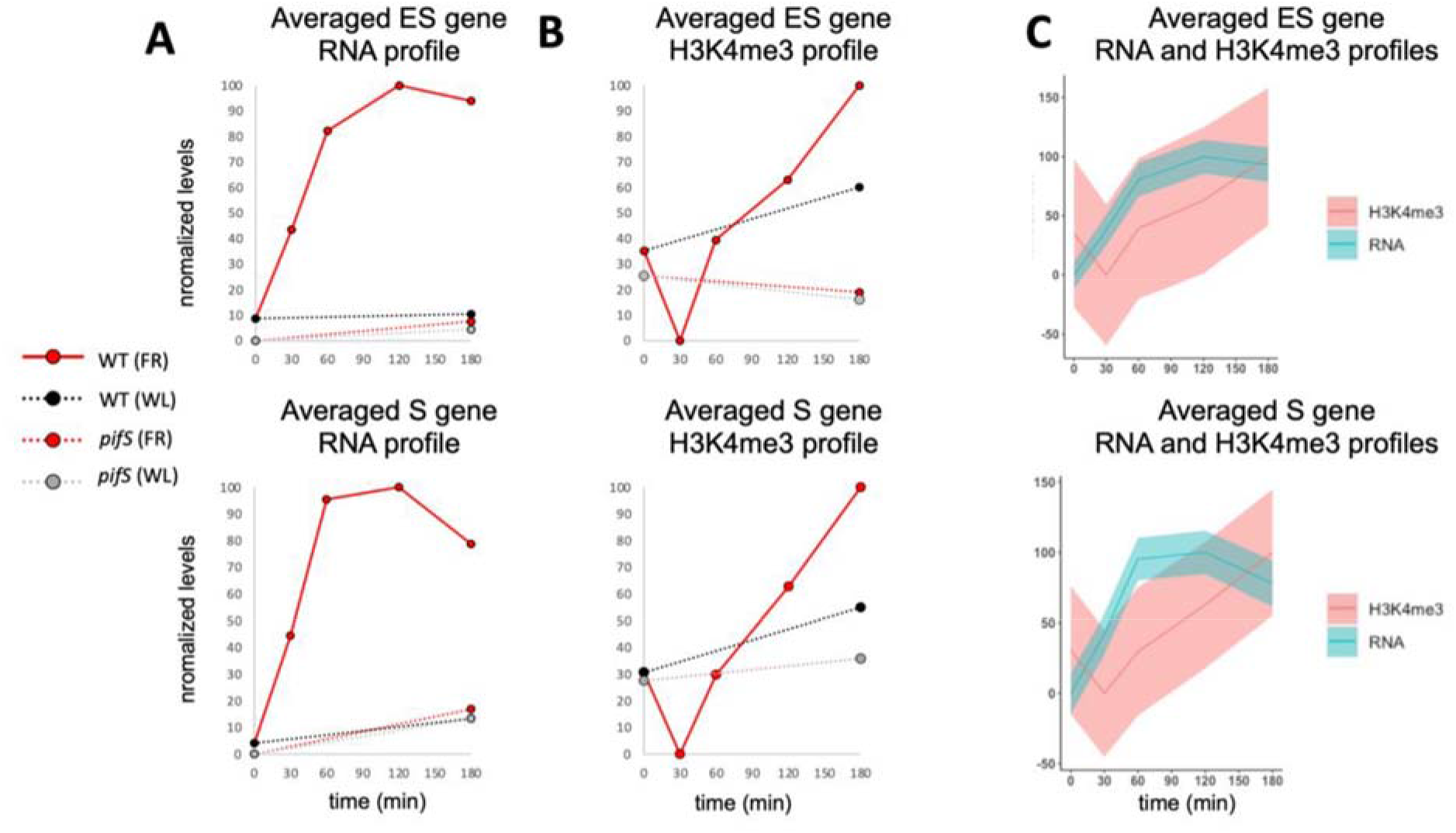
Shade-induced, PIF-dependent increases in transcription precede corresponding increases in H3K4me3 for ES class and S class genes. **A)** Normalized levels of the average transcript profile for all ES class genes (top) or S class genes (bottom) during 180 minute shade-light (FR, red) or white-light (WL, black/gray) treatment for wild-type (WT, solid red/dashed black) or *pifS* (dashed red/dashed gray). **B)**Normalized levels of the average H3K4me3 profile for all ES class genes (top) or S class genes (bottom) during 180 minute shade-light (FR, red) or white-light (WL, black/gray) treatment for wild-type (WT, solid red/dashed black) or *pifS*(dashed red/dashed gray). **C)**Overlays of the RNA (blue) and H3K4me3 (red) profiles from WT seedlings during the FR treatment. Shaded variance indicate normalized SE.

## Discussion

As a prelude to exploring the role of epigenetic factors in light/shade-regulated gene expression, we generated a set of 144 “high-confidence”, PIF-induced DTGs, that we identified by integrating our newly obtained data with previously published analyses. This provided three subclasses of PIF-DTGs, displaying three contrasting patterns of transcriptional responsiveness to light and shade signals (E, ES and S) during young seedling development. By focusing on the shade-responsiveness of these gene sets, we were able to concurrently assess whether differences in the epigenetic landscape might be associated with the observed transcriptional pattern differences, and whether comparison of the temporal patterns of shade-induced transcript and H3K4me3 changes might indicate the potential sequence of such changes.

Broadly speaking our data are consistent with previous studies reporting that high H3K4me3 levels are correlated with actively transcribing genes. However, comparison of our integrated RNA-seq and ChIP-seq analyses over time following shade exposure, showed no clear temporal coincidence of transcript and H3K4me3 levels. On the contrary, for the shade-induced PIF DTGs, we found that, on average, transcript levels rise before their corresponding H3K4me3 levels rise (**Figure 4**). These results indicate that H3K4me3 plays little or no role in causing or priming the rapid, shade-induced transcriptional responsiveness of these genes. Instead, the data are more consistent with previous reports indicating that high levels of transcription from a given locus leads to trimethylation of H3K4 (Le Martelot et al., 2012; Kuang et al., 2014).

Moreover, consideration of our current findings in the context of recent advances in understanding chromatin involvement in controlling plant gene expression, suggests an intriguing possible role for H3K4me3 in shade-regulated expression through the PIF-signaling hub. Willige et al. (Willige et al., 2021) reported that shade rapidly (within 5 minutes) induces the binding of PIF7 to the promoter of the *ATHB2* gene, and similarly rapidly triggers ejection of the histone variant H2A.Z, as well as increasing H3K9 acetylation (H3K9ac). These findings indicate that PIF7 occupancy of target gene promoters can shape the local chromatin status in response to shade. These changes preceded changes in gene expression, leading to the conclusion that chromatin remodeling is not a consequence of transcriptional activation. Given, firstly, that our data indicate, conversely to those of Willige et al. (Willige et al., 2021), that the shade-invoked, PIF-mediated induction of target gene expression appears to precede the increases in H3K4me3 levels at those genes; and secondly, that these H3K4me3 increases are considerably slower than both (a) the shade-induced increases in H3K9ac levels reported by Willige et al. (Willige et al., 2021), and (b) the light-triggered decrease of this mark in dark-grown seedlings observed by González-Grandío (González-Grandío et al., 2022), it appears that H3K4me3 may be a trailing indicator of the expression status of shade-induced genes. This conclusion raises the possibility that H3K4me3 may function to stabilize the active transcriptional state of these genes, thus providing a form of transcriptional memory (Foroozani et al., 2021) as a buffer against exposure to the rapid, random fluctuations between full sunlight and shade that occur within leaf canopies, as a result of breeze-induced movement under natural conditions. The mechanism by which PIF binding activates H3K4 trimethylation remains to be determined.

Collectively, these changes in chromatin landscape add another dimension of complexity to the multilayered network of mechanisms and pathways that regulate and intersect with the phy-PIF module. The phy family have dual photosensory and thermosensory functions, monitoring both light and temperature signals from the environment, that are then transduced through the PIFs (Leivar and Monte, 2014; Legris et al., 2016; Paik et al., 2017). In addition, the PIF family function as a signaling hub for multiple other signaling pathways, that include the core circadian oscillator, via the TOC1 component and its PRR relatives (Soy et al., 2016; Martín et al., 2018; Zhang et al., 2020), the hormones gibberellic acid, abscisic acid, jasmonic acid, ethylene and brassinosteroids (Leivar and Monte, 2014; Paik et al., 2017), as well as interacting with the blue-light photoreceptor, cryptochrome 2 (CRY2) (Más et al., 2000; Pedmale et al., 2016), and numerous other factors, which together are involved in a diversity of molecular functions, that include transcriptional and posttranscriptional modulation (Wang et al., 2021), phosphorylation, ubiquitination, and degradation. Moreover, many of these light-induced interactions appear to take place in nuclear photobodies (Legris et al., 2019), functioning either as a concentrated milieu of dynamically changing, multi-component complexes, driving enhanced intermolecular interactions (Wang et al., 2021), or as foci of sequestration, as shown for PIF7 (Willige et al., 2021).

## Materials and Methods

### Plant growth and phenotyping

All seeds were stratified for 4 days at 4° before germination. Germination was induced by 3h of incubation under 30 μmol m^-2^ s^-1^ WL at 21° followed by a 5 min saturating pulse of FR light. Seedlings were grown for 3 days at 21° in complete darkness or under 30 μmol m^-2^ s^-1^ WL (R/FR = 6-8). For FR light treatment, seedlings were grown for 3 days in WL before exposing them to simulated shade (30 μmol m^-2^ s^-1^, R/FR ~ 0.3). R light was defined as 640-680 nm and FR was defined as 710-750nm.

Hypocotyl measurements were performed on seedlings grown at 23° for 2 days in WL and either exposed to simulated shade for 4 days or kept in constant WL for 4 days. Three independent biological replicates were performed, each of which involved the plating of at least 30 seeds of each genotype all on the same plate. Plates were photographed with a high-resolution camera and hypocotyl lengths were measured via ImageJ. Mean hypocotyl length of each genotype was determined by averaging the means of the three replicates. Standard error was determined by dividing the standard deviation between all three replicas by the square root of 3. Student’s T-test was performed for determination of p-values.

### RNA-seq analysis

RNA was isolated as described (Zhang et al., 2013). Total RNA was extracted from 3-day-old seedlings using a QIAshredder and RNeasy Plus Mini Kit (Qiagen) according to manufacturer’s instructions. RNA libraries for sequencing were prepared at the Functional Genomics Laboratory at UC Berkeley using a KAPA RNA HyperPrep Kit (Roche) according to the manufacturer’s instructions.

RNA libraries were sequenced by the Genomic Sequencing Facility at UC Berkeley. Multiplexed RNA libraries were sequenced by 100-bp paired-end sequencing over two lanes on a HiSeq4000.

For mapping and analysis of RNA-seq experiments, reads were mapped to the Arabidopsis genome (TAIR10) by TopHat (Trapnell et al., 2009) (max intron length = 3000, inner mean distance = 200, inner distance standard deviation = 100, minimal allowed intron size = 25). Assembled reads were counted using featureCounts (Liao et al., 2014) and differential expression was determined via DESeq2 (Love et al., 2014) (log2FC > 1 or log_2_FC < −1; p-val < 0.05).

### Generation of PIF DTG list and subcategorization into E, ES and S classes

To identify PIF DTGs, we first imposed strict statistically-significant two-fold (SSTF) cutoffs and selected all 764 genes whose expression levels decreased in response to red light (Pfeiffer et al., 2014) and/or increased in response to shade-light (this study). We then further narrowed our list to include only those genes that show a dependence on PIFs for their expression by combining the previously published RNA-seq data from the *pifq* mutant grown in darkness (Pfeiffer et al., 2014), with our newly obtained RNA-seq data, obtained using the *pifqpif6pif7* mutant (*pifS*) grown in WL and exposed to 3h shade-light. We selected only those genes that were SSTF induced in WT relative to their levels in the corresponding *pif* mutant. By filtering out those genes that were not among the 764 light-responsive genes identified above, we were left with 278 PIF-dependent, light-responsive genes. Selecting only those genes that were found to be bound by one or more PIF (Hornitschek et al., 2012; Oh et al., 2012; Zhang et al., 2013; Pfeiffer et al., 2014; Chung et al., 2020) yielded 169 genes (**Table 1**).

We subcategorized genes into E, ES and S classes as in Leivar et al, 2012. Class E (formerly Class L) represents genes whose dark-grown wild-type transcript levels are both (a) SSTF higher than those in dark-grown *pifq* and (b) SSTF repressed by the initial red light (R) signal in WT. Although some Class E genes show a degree of re-induction in the shade, this is weaker (*i.e*. non-SSTF), and the PIF-dependency is less, than initially in the dark (**Figure 2**). Conversely, Class S (formerly Class R) represents genes that do display SSTF induction by shade-light, as well as PIF-dependent SSTF induction in the shade, but that do not exhibit a SSTF response to either: (a) the PIFs in dark-grown seedlings, or (b) red light exposure (**Figure 2**). Finally, Class ES (formerly Class M) represents those genes that display SSTF, mutually-converse responsiveness to the onset of the light and shade-light signals, respectively, as well as PIF-dependent SSTF induction, both in the dark and in shade-light (**Figure 2**).

A subset of these E, ES and S class genes exhibited anomalous transcription profiles. We manually removed these 25 genes because they were either highly expressed in WL (6 genes), were induced, rather than repressed, by red light (16 genes), were lowly expressed (1 gene) or were otherwise likely to be artifactual (2 genes). The remaining 144 PIF DTGs were then resorted using relaxed cutoffs. Of the non-anomalous genes first categorized as S class, 38 showed a R-dependent reduction (p < 0.1) in transcript levels but were excluded from the ES class because they did not show a SSTF reduction in dark-grown *pifq* mutant relative to WT. These genes were reclassified as ES. Two E class genes were also reclassified as ES genes because they exhibited a statistically-significant upregulation in response to FR despite not being SSTF downregulated in the *pifS* mutant. Ultimately, we were left with 17 E genes, 56 ES genes and 71 S genes **(Table 1**).

### H3K4me3 ChIP-seq analysis

DNA libraries were sequenced by the Genomic Sequencing Facility at UC Berkeley. The multiplexed DNA libraries were sequenced by 50-bp single-end sequencing over two lanes on a HiSeq4000.

For mapping and analysis of ChIP-seq experiments, reads were mapped to the Arabidopsis genome (TAIR10) by BowTie2 (Langmead and Salzberg, 2012) and uniquely-mapping reads were first used to call peaks using BayesPeak (Spyrou et al., 2009) (Bioconductor 3.6; binsize = 300, peaks with a PP>0.999 in all 3 biological replicates) or MACS (Zhang et al., 2008). H3K4me3 peaks calculated using BayesPeak and MACS2 could only be unambiguously assigned to the transcriptional start sites (TSS) of 102 of the 144 E, ES, and S class genes. To ensure consistency in analysis, we therefore manually assigned peaks to all of the PIF DTGs by creating 300bp windows centered on the TSS.

To quantify the H3K4me3 peaks and measure differences between time points we used DiffBind (Ross-Innes et al., 2012) and DESeq2 (Love et al., 2014). Because the changes in magnitude of H3K4me3 levels were far smaller than the changes in transcript abundance, we used DESeq2 to calculate variance-stabilizing transformations (VSTs) across the time course experiment for both H3K4me3 levels and transcript levels. This enabled comparison of relative changes in H3K4me3 levels to the corresponding changes in transcription for a given gene or class of genes.

## Supplemental Data

**Supplemental Table S1.** List of PIF-induced DTGs identified in this study and whether or not they have been previously identified as a PIF DTG.

## Acknowledgements

We thank Eduardo González-Grandío and James Tepperman for extensive discussions, critical feedback and for facilitating an excellent research environment. We also thank Alexander Vergara and Martí Quevedo for assistance in analyzing ChIP data and Luis Cervela-Cardona for critical reading of the manuscript.

## Author Contributions

RHC and PHQ designed the research. RHC, JD and YZ performed research. RHC, JD, YZ and PHQ analyzed data. RHC and PHQ wrote the paper.

## Parsed Citations

Al-Sady B, Kikis EA, Monte E, Quail PH (2008) Mechanistic duality of transcription factor function in phytochrome signaling. Proc Natl Acad Sci U S A 105: 2232–2237

Bernstein BE, Humphrey EL, Erlich RL, Schneider R, Bouman P, Liu JS, Kouzarides T, Schreiber SL (2002) Methylation of histone H3 Lys 4 in coding regions of active genes. Proc Natl Acad Sci U S A 99: 8695–8700

Bourbousse C, Ahmed I, Roudier F, Zabulon G, Blondet E, Balzergue S, Colot V, Bowler C, Barneche F (2012) Histone H2B monoubiquitination facilitates the rapid modulation of gene expression during Arabidopsis photomorphogenesis. PLoS Genet 8: e1002825

Bourbousse C, Barneche F, Laloi C (2019) Plant Chromatin Catches the Sun. Front Plant Sci 10: 1728

Charron JB, He H, Elling AA, Deng XW (2009) Dynamic landscapes of four histone modifications during deetiolation in Arabidopsis. Plant Cell 21: 3732–3748

Chung BYW, Balcerowicz M, Di Antonio M, Jaeger KE, Geng F, Franaszek K, Marriott P, Brierley I, Firth AE, Wigge PA (2020) An RNA thermoswitch regulates daytime growth in Arabidopsis. Nat Plants 6: 522–532

Dalton JC, Bätz U, Liu J, Curie GL, Quail PH (2016) A Modified Reverse One-Hybrid Screen Identifies Transcriptional Activation Domains in PHYTOCHROME-INTERACTING FACTOR 3. Front Plant Sci 7: 881

de Lucas M, Davière JM, Rodríguez-Falcón M, Pontin M, Iglesias-Pedraz JM, Lorrain S, Fankhauser C, Blázquez MA, Titarenko E, Prat S (2008) A molecular framework for light and gibberellin control of cell elongation. Nature 451: 480–484

de Wit M, Ljung K, Fankhauser C (2015) Contrasting growth responses in lamina and petiole during neighbor detection depend on differential auxin responsiveness rather than different auxin levels. New Phytol 208: 198–209

Ding Y, Ndamukong I, Xu Z, Lapko H, Fromm M, Avramova Z (2012) ATX1-generated H3K4me3 is required for efficient elongation of transcription, not initiation, at ATX1-regulated genes. PLoS Genet 8: e1003111

Fiorucci AS, Bourbousse C, Concia L, Rougée M, Deton-Cabanillas AF, Zabulon G, Layat E, Latrasse D, Kim SK, Chaumont N, Lombard B, Stroebel D, Lemoine S, Mohammad A, Blugeon C, Loew D, Bailly C, Bowler C, Benhamed M, Barneche F (2019) Arabidopsis S2Lb links AtCOMPASS-like and SDG2 activity in H3K4me3 independently from histone H2B monoubiquitination. Genome Biol 20: 100

Fisher AJ, Franklin KA (2011) Chromatin remodelling in plant light signalling. Physiol Plant 142: 305–313

Foroozani M, Vandal MP, Smith AP (2021) H3K4 trimethylation dynamics impact diverse developmental and environmental responses in plants. Planta 253: 4

González-Grandío E, Álamos S, Zhang Y, Dalton-Roesler J, Niyogi KK, García HG, Quail PH (2022) Chromatin Changes in Phytochrome Interacting Factor-Regulated Genes Parallel Their Rapid Transcriptional Response to Light. Front Plant Sci 13: 803441

Hornitschek P, Kohnen MV, Lorrain S, Rougemont J, Ljung K, López-Vidriero I, Franco-Zorrilla JM, Solano R, Trevisan M, Pradervand S, Xenarios I, Fankhauser C (2012) Phytochrome interacting factors 4 and 5 control seedling growth in changing light conditions by directly controlling auxin signaling. Plant J 71: 699–711

Huq E, Al-Sady B, Hudson M, Kim C, Apel K, Quail PH (2004) Phytochrome-interacting factor 1 is a critical bHLH regulator of chlorophyll biosynthesis. Science 305: 1937–1941

Huq E, Quail PH (2002) PIF4, a phytochrome-interacting bHLH factor, functions as a negative regulator of phytochrome B signaling in Arabidopsis. EMBO J 21: 2441–2450

Khanna R, Huq E, Kikis EA, Al-Sady B, Lanzatella C, Quail PH (2004) A novel molecular recognition motif necessary for targeting photoactivated phytochrome signaling to specific basic helix-loop-helix transcription factors. Plant Cell 16: 3033–3044

Kuang Z, Cai L, Zhang X, Ji H, Tu BP, Boeke JD (2014) High-temporal-resolution view of transcription and chromatin states across distinct metabolic states in budding yeast. Nat Struct Mol Biol 21: 854–863

Langmead B, Salzberg SL (2012) Fast gapped-read alignment with Bowtie 2. Nat Methods 9: 357–359

Le Martelot G, Canella D, Symul L, Migliavacca E, Gilardi F, Liechti R, Martin O, Harshman K, Delorenzi M, Desvergne B, Herr W, Deplancke B, Schibler U, Rougemont J, Guex N, Hernandez N, Naef F, Consortium C (2012) Genome-wide RNA polymerase II profiles and RNA accumulation reveal kinetics of transcription and associated epigenetic changes during diurnal cycles. PLoS Biol 10: e1001442

Legris M, Ince Y, Fankhauser C (2019) Molecular mechanisms underlying phytochrome-controlled morphogenesis in plants. Nat Commun 10: 5219

Legris M, Klose C, Burgie ES, Rojas CC, Neme M, Hiltbrunner A, Wigge PA, Schäfer E, Vierstra RD, Casal JJ (2016) Phytochrome B integrates light and temperature signals in Arabidopsis. Science 354: 897–900

Leivar P, Monte E (2014) PIFs: systems integrators in plant development. Plant Cell 26: 56–78

Leivar P, Monte E, Al-Sady B, Carle C, Storer A, Alonso JM, Ecker JR, Quail PH (2008) The Arabidopsis phytochrome-interacting factor PIF7, together with PIF3 and PIF4, regulates responses to prolonged red light by modulating phyB levels. Plant Cell 20: 337–352

Leivar P, Monte E, Oka Y, Liu T, Carle C, Castillon A, Huq E, Quail PH (2008) Multiple phytochrome-interacting bHLH transcription factors repress premature seedling photomorphogenesis in darkness. Curr Biol 18: 1815–1823

Leivar P, Quail PH (2011) PIFs: pivotal components in a cellular signaling hub. Trends Plant Sci 16: 19–28

Leivar P, Tepperman JM, Cohn MM, Monte E, Al-Sady B, Erickson E, Quail PH (2012) Dynamic antagonism between phytochromes and PIF family basic helix-loop-helix factors induces selective reciprocal responses to light and shade in a rapidly responsive transcriptional network in Arabidopsis. Plant Cell 24: 1398–1419

Leivar P, Tepperman JM, Monte E, Calderon RH, Liu TL, Quail PH (2009) Definition of early transcriptional circuitry involved in light-induced reversal of PIF-imposed repression of photomorphogenesis in young Arabidopsis seedlings. Plant Cell 21: 3535–3553

Li L, Ljung K, Breton G, Schmitz RJ, Pruneda-Paz J, Cowing-Zitron C, Cole BJ, Ivans LJ, Pedmale UV, Jung HS, Ecker JR, Kay SA, Chory J (2012) Linking photoreceptor excitation to changes in plant architecture. Genes Dev 26: 785–790

Liao Y, Smyth GK, Shi W (2014) featureCounts: an efficient general purpose program for assigning sequence reads to genomic features. Bioinformatics 30: 923–930

Liu N, Fromm M, Avramova Z (2014) H3K27me3 and H3K4me3 chromatin environment at super-induced dehydration stress memory genes of Arabidopsis thaliana. Mol Plant 7: 502–513

Liu X, Chen CY, Wang KC, Luo M, Tai R, Yuan L, Zhao M, Yang S, Tian G, Cui Y, Hsieh HL, Wu K (2013) PHYTOCHROME INTERACTING FACTOR3 associates with the histone deacetylase HDA15 in repression of chlorophyll biosynthesis and photosynthesis in etiolated Arabidopsis seedlings. Plant Cell 25: 1258–1273

Love MI, Huber W, Anders S (2014) Moderated estimation of fold change and dispersion for RNA-seq data with DESeq2. Genome Biol 15: 550

Martín G, Rovira A, Veciana N, Soy J, Toledo-Ortiz G, Gommers CMM, Boix M, Henriques R, Minguet EG, Alabadí D, Halliday KJ, Leivar P, Monte E (2018) Circadian Waves of Transcriptional Repression Shape PIF-Regulated Photoperiod-Responsive Growth in Arabidopsis. Curr Biol 28: 311–318.e315

Martínez-García JF, Moreno-Romero J (2020) Shedding light on the chromatin changes that modulate shade responses. Physiol Plant 169: 407–417

Mizuno T, Oka H, Yoshimura F, Ishida K, Yamashino T (2015) Insight into the mechanism of end-of-day far-red light (EODFR)-induced shade avoidance responses in Arabidopsis thaliana. Biosci Biotechnol Biochem 79: 1987–1994

Monte E, Tepperman JM, Al-Sady B, Kaczorowski KA, Alonso JM, Ecker JR, Li X, Zhang Y, Quail PH (2004) The phytochromeinteracting transcription factor, PIF3, acts early, selectively, and positively in light-induced chloroplast development. Proc Natl Acad Sci U S A 101: 16091–16098

Más P, Devlin PF, Panda S, Kay SA (2000) Functional interaction of phytochrome B and cryptochrome 2. Nature 408: 207–211

Ni M, Tepperman JM, Quail PH (1998) PIF3, a phytochrome-interacting factor necessary for normal photoinduced signal transduction, is a novel basic helix-loop-helix protein. Cell 95: 657–667

Oh E, Zhu JY, Wang ZY (2012) Interaction between BZR1 and PIF4 integrates brassinosteroid and environmental responses. Nat Cell Biol 14: 802–809

Paik I, Kathare PK, Kim JI, Huq E (2017) Expanding Roles of PIFs in Signal Integration from Multiple Processes. Mol Plant 10: 1035–1046

Pedmale UV, Huang SC, Zander M, Cole BJ, Hetzel J, Ljung K, Reis PAB, Sridevi P, Nito K, Nery JR, Ecker JR, Chory J (2016) Cryptochromes Interact Directly with PIFs to Control Plant Growth in Limiting Blue Light. Cell 164: 233–245

Penfield S, Josse EM, Halliday KJ (2010) A role for an alternative splice variant of PIF6 in the control of Arabidopsis primary seed dormancy. Plant Mol Biol 73: 89–95

Perrella G, Kaiserli E (2016) Light behind the curtain: photoregulation of nuclear architecture and chromatin dynamics in plants. New Phytol 212: 908–919

Pfeiffer A, Shi H, Tepperman JM, Zhang Y, Quail PH (2014) Combinatorial complexity in a transcriptionally centered signaling hub in Arabidopsis. Mol Plant 7: 1598–1618

Pham VN, Kathare PK, Huq E (2018) Phytochromes and Phytochrome Interacting Factors. Plant Physiol 176: 1025–1038

Quail PH, Boylan MT, Parks BM, Short TW, Xu Y, Wagner D (1995) Phytochromes: photosensory perception and signal transduction. Science 268: 675–680

Ross-Innes CS, Stark R, Teschendorff AE, Holmes KA, Ali HR, Dunning MJ, Brown GD, Gojis O, Ellis IO, Green AR, Ali S, Chin SF, Palmieri C, Caldas C, Carroll JS (2012) Differential oestrogen receptor binding is associated with clinical outcome in breast cancer. Nature 481: 389–393

Sakamoto K, Nagatani A (1996) Nuclear localization activity of phytochrome B. Plant J 10: 859–868\

Soy J, Leivar P, González-Schain N, Martín G, Diaz C, Sentandreu M, Al-Sady B, Quail PH, Monte E (2016) Molecular convergence of clock and photosensory pathways through PIF3-TOC1 interaction and co-occupancy of target promoters. Proc Natl Acad Sci U S A 113: 4870–4875

Spyrou C, Stark R, Lynch AG, Tavaré S (2009) BayesPeak: Bayesian analysis of ChIP-seq data. BMC Bioinformatics 10: 299

Tepperman JM, Zhu T, Chang HS, Wang X, Quail PH (2001) Multiple transcription-factor genes are early targets of phytochrome A signaling. Proc Natl Acad Sci U S A 98: 9437–9442

Trapnell C, Pachter L, Salzberg SL (2009) TopHat: discovering splice junctions with RNA-Seq. Bioinformatics 25: 1105–1111

Wang X, Jiang B, Gu L, Chen Y, Mora M, Zhu M, Noory E, Wang Q, Lin C (2021) A photoregulatory mechanism of the circadian clock in Arabidopsis. Nat Plants 7: 1397–1408

Willige BC, Zander M, Yoo CY, Phan A, Garza RM, Trigg SA, He Y, Nery JR, Chen H, Chen M, Ecker JR, Chory J (2021) PHYTOCHROME-INTERACTING FACTORs trigger environmentally responsive chromatin dynamics in plants. Nat Genet 53: 955–961

Zhang X, Bernatavichute YV, Cokus S, Pellegrini M, Jacobsen SE (2009) Genome-wide analysis of mono-, di-and trimethylation of histone H3 lysine 4 in Arabidopsis thaliana. Genome Biol 10: R62

Zhang Y, Liu T, Meyer CA, Eeckhoute J, Johnson DS, Bernstein BE, Nusbaum C, Myers RM, Brown M, Li W, Liu XS (2008) Modelbased analysis of ChlP-Seq (MACS). Genome Biol 9: R137

Zhang Y, Mayba O, Pfeiffer A, Shi H, Tepperman JM, Speed TP, Quail PH (2013) A quartet of PIF bHLH factors provides a transcriptionally centered signaling hub that regulates seedling morphogenesis through differential expression-patterning of shared target genes in Arabidopsis. PLoS Genet 9: e1003244

Zhang Y, Pfeiffer A, Tepperman JM, Dalton-Roesler J, Leivar P, Gonzalez Grandio E, Quail PH (2020) Central clock components modulate plant shade avoidance by directly repressing transcriptional activation activity of PIF proteins. Proc Natl Acad Sci U S A 117: 3261–3269

