## Supplemental Table 1 for "Shade-induced transcription of PIF-Direct-Target Genes precedes H3K4-trimethylation chromatin modification rises"

**Supplemental Table 1**: **List of PIF-induced DTGs identified in this study and whether or not they have been previously identified as a PIF DTG.**

| locus | name | Final class | Oh 2012 | Zhang 2013 | Leivar 2014 | Nozue 2011 | Shin 2009 | Pfieffer 2014 | Hornitschek 2012 | Martin 2016 | Chung 2020 | Other | previously described PIF DTG |
|---|---|---|---|---|---|---|---|---|---|---|---|---|---|
| AT1G29460 | SAUR65 | S | X | | X | | | | X | X | X | 1 | yes |
| AT1G25560 | EDF1 | ES | | | | | | | | | | | no |
| AT1G54120 | At1g54120 | ES | | | | | | | | | | | no |
| AT3G05640 | EGR1 | ES | | | | | | | | | | | no |
| AT3G62070 | At3g62070 | ES | | | | | | | | | | | no |
| AT1G29430 | SAUR62 | ES | | | | | | | | | | | no |
| AT1G29465 | At1g29465 | S | | | | | | | | | | | no |
| AT1G49780 | PUB26 | S | | | | | | | | | | | no |
| AT4G32290 | At4g32290 | S | | | | | | | | | | | no |
| AT5G43890 | YUC5 | S | | | | | | | | | | 2 | yes |
| AT3G62100 | IAA30 | S | | | | | | | | | | | no |
| AT4G24275 | At4g24275 | S | | | | | | | | | | | no |
| AT2G45420 | LBD18 | S | | | | | | | | | | | no |
| AT3G44310 | NIT1 | S | | | | | | | | | | 3 | yes |
| AT5G16200 | At5g16200 | S | | | | | | | | | | | no |
| AT1G52565 | At1g52565 | S | | | | | | | | | | | no |
| AT3G50350 | At3g50350 | S | | | | | | | | | | | no |
| AT5G02260 | EXP9 | E | X | | X | | X | X | | X | | | yes |
| AT5G02580 | At5g02580 | E | X | | X | | X | X | | X | | | yes |
| AT5G67020 | At5g67020 | E | X | | | | | | | | | | yes |
| AT5G02190 | PCS1 | E | X | | | | | X | | X | X | | yes |
| AT1G07090 | LSH6 | E | X | X | X | X | | X | X | X | | | yes |
| AT1G67265 | DVL3 | E | X | | | | | X | | X | | | yes |
| AT1G60060 | At1g60060 | E | X | | X | X | X | | | X | | | yes |
| AT5G15830 | BZIP3 | E | | | X | | | | | X | | | yes |
| AT4G37740 | GRF2 | E | X | X | X | | X | X | | X | | | yes |
| AT5G50175 | At5g50175 | E | | | | | | X | | X | | | yes |
| AT4G36010 | At4g36010 | E | X | X | X | | | X | | X | | | yes |
| AT4G10020 | HSD5 | E | X | X | X | | X | X | | X | | | yes |
| AT3G25730 | EDF3 | E | X | | X | X | X | | | X | | | yes |
| AT3G28340 | GATL10 | E | X | | X | X | X | X | | X | | | yes |
| AT1G58410 | At1g58410 | E | | X | | | X | | | X | | | yes |
| AT3G53200 | MYB27 | E | X | | | | | X | | X | | | yes |
| AT5G53980 | ATHB52 | E | X | X | X | | X | X | X | X | | | yes |
| AT2G42870 | PAR1 | ES | X | | X | | | | X | | | | yes |
| AT3G59900 | ARGOS | ES | X | X | X | | | | X | X | | | yes |
| AT2G44910 | ATHB-4 | ES | X | | X | | | | X | | X | | yes |
| AT5G28300 | GT2L | ES | X | X | X | | | | X | X | | | yes |
| AT1G13260 | RAV1 | ES | X | | X | | | | | | | | yes |
| AT5G44260 | TZF5 | ES | X | | X | | | | X | | X | | yes |
| AT5G02760 | APD7 | ES | | | | | | X | X | | | | yes |
| AT5G62280 | At5g62280 | ES | | | X | X | | | X | | X | | yes |
| AT5G46330 | FLS2 | ES | | X | | | | | X | | X | | yes |
| AT2G44080 | ARL | ES | X | | | | | | | X | | | yes |
| AT3G60390 | HAT3 | ES | X | | X | | | | | | | | yes |
| AT5G25190 | ESE3 | ES | X | | X | X | | | X | | | | yes |
| AT3G60520 | At3g60520 | ES | X | | X | | | | | | | | yes |
| AT4G28240 | BGL1 | ES | | | X | | | | X | | | | yes |
| AT2G45210 | SAUR36 | ES | X | X | X | X | | X | | X | | | yes |
| AT2G43060 | IBH1 | ES | X | | X | X | X | X | X | X | | | yes |
| AT5G02540 | At5g02540 | ES | X | | X | | X | X | X | X | X | | yes |
| AT3G15540 | IAA19 | ES | X | | X | | | X | X | X | | | yes |
| AT3G21330 | At3g21330 | ES | | | | | X | X | | | | | yes |
| AT5G63650 | SNRK2.5 | ES | X | | X | | X | X | X | X | | | yes |
| AT5G07010 | ST2A | ES | X | | X | | X | X | X | X | X | | yes |
| AT5G01790 | At5g01790 | ES | X | | X | X | | X | | X | | | yes |
| AT1G10550 | XTH33 | ES | X | | X | X | | X | | X | X | | yes |
| AT3G61830 | ARF18 | ES | | | X | X | X | X | | X | | | yes |
| AT4G16780 | ATHB-2 | ES | X | X | X | | X | X | X | X | X | | yes |

| | | | | | | | | | | | | |
|---|---|---|---|---|---|---|---|---|---|---|---|---|
| AT4G35720 | At4g35720 | ES | X | | X | | X | X | | X | | yes |
| AT4G14130 | XTR7 | ES | X | | X | | | X | | X | | yes |
| AT4G32280 | IAA29 | ES | X | | X | | X | X | X | | | yes |
| AT5G65800 | ACS5 | ES | | | X | | X | X | | X | | yes |
| AT4G31380 | FLP1 | ES | X | | | | X | | | X | X | yes |
| AT2G46970 | PIL1 | ES | X | | | | X | X | | X | | yes |
| AT5G05965 | At5g05965 | ES | X | | | | X | X | | X | | yes |
| AT5G09970 | CYP78A7 | ES | X | | | | | | | | | yes |
| AT1G21050 | At1g21050 | ES | X | | X | | | | | | | yes |
| AT5G59010 | BSK1 | ES | X | X | | | | | X | X | | yes |
| AT3G61460 | BRH1 | ES | | | X | | | | X | | X | yes |
| AT1G21830 | At1g21830 | ES | | | X | | | | | | | yes |
| AT3G50340 | At3g50340 | ES | | | X | | | | | | | yes |
| AT4G22780 | ACR7 | ES | X | | X | X | | | X | | | yes |
| AT2G28400 | At2g28400 | ES | | | X | | | | | X | | yes |
| AT4G25260 | PMEI7 | ES | X | | X | | X | X | X | X | | yes |
| AT5G46240 | KAT1 | ES | X | | X | | | | X | | | yes |
| AT5G18030 | SAUR21 | ES | X | | | | | | | X | X | yes |
| AT4G38860 | SAUR16 | ES | X | | | | | | | | | yes |
| AT1G02400 | GA2OX6 | ES | X | | X | | | | | | | yes |
| AT1G75450 | CKX5 | ES | X | | X | X | X | X | | | X | yes |
| AT5G18060 | SAUR23 | ES | X | | X | X | | X | X | X | X | yes |
| AT4G37770 | ACS8 | ES | X | X | X | X | | | X | | | yes |
| AT4G13790 | SAUR25 | ES | X | X | X | | X | X | | X | | yes |
| AT3G62090 | PIF6 | ES | X | | X | X | X | X | | X | | yes |
| AT3G12820 | MYB10 | ES | X | X | X | X | X | | | X | | yes |
| AT3G21320 | At3g21320 | S | X | | | | | | | | | yes |
| AT5G22500 | FAR1 | S | X | | X | | | | X | X | | yes |
| AT4G28720 | YUC8 | S | X | X | X | | | | X | | X | yes |
| AT1G04180 | YUC9 | S | | | X | | | | | | | yes |
| AT5G18050 | SAUR22 | S | X | | | | | | | X | X | yes |
| AT1G02350 | At1g02350 | S | X | | | | | | | | | yes |
| AT5G66080 | APD9 | S | X | | X | | | | X | X | X | yes |
| AT3G23030 | IAA2 | S | X | | X | | | | X | | | yes |
| AT5G47370 | HAT2 | S | | | X | | | | X | | | yes |
| AT4G14560 | IAA1 | S | | | X | | | | | | | yes |
| AT5G25460 | DGR2 | S | X | | | | | | X | | X | yes |
| AT1G36940 | At1g36940 | S | X | | X | | | | | | | yes |
| AT3G23050 | IAA7 | S | | | | | | | X | X | | yes |
| AT2G23170 | GH3.3 | S | | | X | | | | | | | yes |
| AT5G12050 | BG1 | S | | X | X | | | | X | | X | yes |
| AT1G76610 | At1g76610 | S | | | | | | | | | X | yes |
| AT1G75500 | WAT1 | S | X | | | | | | X | | X | yes |
| AT1G21980 | PIP5K1 | S | X | | | | | | | X | | yes |
| AT1G31880 | BRX | S | X | | | | | | | | | yes |
| AT4G39800 | MIPS1 | S | | | | | | | | | | yes |
| AT1G67900 | At1g67900 | S | X | | | | | | | | X | yes |
| AT5G16023 | DVL1 | S | X | | | | | | | | | yes |
| AT5G39860 | PRE1 | S | X | | X | X | | | X | | X | yes |
| AT4G34760 | SAUR50 | S | X | | | | | | | | | yes |
| AT4G37390 | GH3.2 | S | | | | | | X | | | | yes |
| AT1G75490 | At1g75490 | S | X | | | | | X | | | | yes |
| AT4G27280 | CMI1 | S | | | X | | | | | | | yes |
| AT4G27310 | BBX28 | S | X | | | | X | | | | | yes |
| AT5G59220 | HAI1 | S | X | | | | | X | | | | yes |
| AT1G60190 | PUB19 | S | X | | | | | | X | | | yes |
| AT2G40610 | EXP8 | S | X | | X | X | | | X | | | yes |
| AT5G54510 | GH3.6 | S | X | | | | | | X | | | yes |
| AT5G60840 | At5g60840 | S | X | | X | | | | | | | yes |
| AT4G09890 | At4g09890 | S | X | | X | X | | | X | | | yes |
| AT5G62220 | GT18 | S | X | | | | | | X | | | yes |
| AT3G19380 | PUB25 | S | | | | | | | X | | | yes |
| AT1G21910 | DREB26 | S | X | | | | | | | | | yes |
| AT3G03850 | SAUR26 | S | X | | | | | | | | | yes |

| | | | | | | | | | | | | |
|---|---|---|---|---|---|---|---|---|---|---|---|---|
| AT3G03840 | SAUR27 | S | X | | | | | | X | | | yes |
| AT2G18010 | SAUR10 | S | X | | | | | | | | | yes |
| AT3G03830 | SAUR28 | S | X | X | X | | | | X | X | | yes |
| AT4G34770 | SAUR1 | S | X | X | X | | X | | X | X | | yes |
| AT1G29500 | SAUR66 | S | X | | X | X | | | X | | | yes |
| AT3G55840 | At3g55840 | S | X | | X | | | | | | | yes |
| AT5G18020 | SAUR20 | S | X | | | | | | | X | X | yes |
| AT1G29440 | SAUR63 | S | X | | X | | | | X | | | yes |
| AT1G29450 | SAUR64 | S | X | | | | | | X | | | yes |
| AT1G69160 | WIP1 | S | X | | | | | X | X | | | yes |
| AT5G18010 | SAUR19 | S | X | | | | | X | | X | | yes |
| AT3G50800 | At3g50800 | S | X | | | | | X | | | X | yes |
| AT1G04240 | IAA3 | S | | | | | | X | | | X | yes |
| AT1G76240 | At1g76240 | S | X | X | X | | X | X | X | X | | yes |
| AT1G18400 | BEE1 | S | | | X | | | X | X | X | | yes |
| AT5G66580 | At5g66580 | S | X | | | | | X | | X | X | yes |
| AT3G28857 | PRE5 | S | X | X | | | X | X | | | | yes |
| AT2G14960 | GH3.1 | S | | | | | | X | | | | yes |
| AT1G06080 | ADS1 | S | X | | X | | X | | X | X | | yes |
| AT5G66590 | At5g66590 | S | X | X | X | | X | X | X | X | | yes |
| AT1G16850 | At1g16850 | S | X | | X | | X | X | | X | | yes |


| | | |
|---|---|---|
| Oh 2012 | Oh et al, 2012, Nat Cell Biol | (https://doi.org/10.1038/ncb2545) |
| Zhang 2013 | Zhang et al 2013, PLOS Genetics | (https://doi.org/10.1371/journal.pgen.1003244) |
| Leivar 2014 | Leivar and Monte, 2014, Plant Cell | (https://doi.org/10.1105/tpc.113.120857) |
| Nozue 2011 | Nozue et al, 2011, Plant Physiol | (https://doi.org/10.1104/pp.111.172684) |
| Shin 2009 | Shin et al, 2009, PNAS | (https://doi.org/10.1073/pnas.0812219106) |
| Pfieffer 2014 | Pfieffer et al, 2014, Mol Plant | (https://doi.org/10.1093/mp/ssu087) |
| Hornitschek 2012 | Hornitschek et al, 2012, Plant J | (https://doi.org/10.1111/j.1365-313X.2012.05033.x) |
| Martin 2016 | Martin et al, 2016, Nat Commun | (https://doi.org/10.1038/ncomms11431) |
| Chung 2020 | Chung et al, 2020, Nat Plants | (https://doi.org/10.1038/s41477-020-0633-3) |

1= Sun et al, 2016, PNAS (https://doi.org/10.1073/pnas.1604782113)
2=Goyal et al., 2016, Curr Biol (https://doi.org/10.1016/j.cub.2016.10.001)
3=Oh et al, 2009, Plant Cell (https://doi.org/10.1105/tpc.108.064691)
